# Radiofrequency Remote Control of Thermolysin Activity

**DOI:** 10.1101/781138

**Authors:** Christian B. Collins, Christopher J. Ackerson

## Abstract

Nearly all biological processes are regulated by enzymes, precise control over specific enzymes could create the potential for controlling cellular processes remotely. We have successfully shown that the thermophilic enzyme thermolysin can be remotely activated in 17.76 MHz radiofrequency (RF) fields when covalently attached to 6.1 nm gold coated magnetite nanoparticles. Without raising the bulk solution temperature, we observe enzyme activity as if the solution was 16 ± 2 °C warmer in RF fields, or an increase in enzymatic rate of 129 ± 8%. Kinetics studies show that the activity increase of the enzyme is consistent with the induced fit of a hot enzyme with cold substrate.

## Introduction

The use of temperature changes to modulate biological processes is ubiquitous. Well known examples include cooking to neutralize foodborne pathogens, the polymerase chain reaction (PCR) [1–5], thermal cancer treatment [6–10], and thermal deactivation of enzymes (e.g., restriction endonucleases) [11–13]. These processes all rely on bulk(macro)-scale heating.

The routine use of thermal manipulation of biological processes is ‘bulk heating’ – heating the entire sample. However, the possibility of heating discrete and defined local regions within a sample offers the potential to extend the paradigm of thermal manipulation of biological samples to allow targeting of discrete molecules with minimal collateral heating.

The advent of nanoparticles that convert external stimuli to heat gives rise to the possibility that thermal modulation of biological processes could be spatially targeted with high resolution. Indeed, inorganic particles that heat upon thermal stimuli have been used to modulate biological activity. Gold nanoparticles conjugated to DNA can thermally modulate DNA hybridization [14]. Micron-sized ferromagnetic iron-oxides can thermally control the activity of amylase under 0.34 MHz RF irradiation [15]. Dehalogenase enzymes immobilized in a gel matrix with iron-oxide particles of unknown size could be thermally controlled using the nanoparticles to heat the entire hydrogel [16]. Gold nanorods that heat upon NIR, 800 nm laser stimulation can modulate the activity of glucokinase [17].

Prior examples of particle mediated influence of enzyme activity are done with particles of widely varying size, from micron sized iron-oxides to 30 × 10 nm nanorod [17]. The different sizes of particle imply different spatial resolutions to which heat can be delivered. For instance, micron [18] and 100+ nm sized particles [19] could be effective at cellular resolution—heating nearby cells but not far away cells, while sub-10 nm sized particles may heat at molecular resolution—heating nearby molecules but not far-away molecules. We choose to utilize smaller particles than previously reported, in this sub-10 nm size regime (6.1 nm) to improve heat resolution and increase the surface area to volume ratio, allowing for increased area for heating targets and heat transfer. Figure S1 summarizes the differences in surface area to volume ratio and shows the striking difference for a fixed volume fraction of particles.

Herein, we demonstrate the first thermal manipulation of enzymatic activity by ‘hot’ nanoparticles at molecular resolution. We use 6.1 nm gold coated magnetite particles *covalently linked* to the thermostable protease, thermolysin. The superparamagnetic 4.5 nm magnetite cores heat under radiofrequency (RF) irradiation at 17.76 MHz, as the magnetic field torques the magnetic moments and the particles relax via Neel relaxation mechanisms [20–24].

We observe an increase of thermolysin activity that is proportional to the applied RF power (and B field strength), without increasing the bulk temperature of the solution. The effective temperature of thermolysin can be correlated to different power inputs. We observe unique enzyme kinetics, consistent with a hot enzyme and cold substrate.

## Materials and Methods

### Magnetite Fe_3_O_4_ Core synthesis procedure

Magnetite cores were synthesized by thermal decomposition in diphenyl ether following a published method [25]. Briefly, in a 100 mL 3 neck round bottom flask; 10 mmol 1,2 hexadecane diol and 2 mmol iron (III) acytlacetonate was mixed in 20 mL of diphenyl ether. The solution was then purged and flushed with argon. Then 6 mmol oleylamine (90%) and 6 mmol oleic acid was added via syringe. The solution was then heated to 200°C for 30 minutes to boil off impurities. Then it was heated to 265°C and refluxed for 1 hour then cooled to 30°C, and transferred into scintillation vials for temporary storage. See SI S4 for complete details.

### Gold Coating Procedure

The magnetite cores were coated using published procedure [26]. Briefly, 5 mL of the magnetite cores solution from above, without any rinsing or work up was added to a 100 mL, 2 neck, round bottom flask. To that solution, 1.1 mmol gold (III) acetate, 6 mmol of 1,2-hexadecanediol, 0.75 mmol oleic acid and 3 mmol oleylamine was added in 15 mL diphenyl ether. This solution was stirred vigorously at 35°C while purging with argon for 20 minutes. Under argon and vigorous stirring the solution was then it was heated to 185°C and held at that temperature 1.5 hours. The particles were cleaned up by rinsing 3× with 1:1 mixture of hexanes and ethanol. See SI S5 for complete details.

### Phase Transfer

The phase transfer step proved to be very challenging and a novel two step procedure was used; first an initial exchange and phase transfer from oleylamine coating to 3-mercaptopropanoic acid (3-MPA). Then an exchange of 3-MPA for 11-mercaptoundecanoic acid (11-MUA) to increase solubility in water and biological relevant buffers. In a 20 mL scintillation vial, 2 mL of 17 mg/mL particle solution in heptanes was added with 10 mL of heptanes 5mL of 3-MPA. This is then vortexed overnight. The particles were then precipitated and washed with 5 mL acetone 3×, until no 3-MPA smell could be detected. The particle pellet was then suspended in 1.5 mL 75 mM tetramethylammonium hydroxide (TMAOH) solution to give a red solution [27].

To improve the stability of the particles in a wide range of solutions the 3-MPA ligands were further exchanged for longer 11-MUA ligands. To the 1.5 mL batch of phase transferred particles, 3.4 mg of 11-MUA was added and the mixture was sonicated for 20 minutes to facilitate dissolution of the 11-MUA. The mixture was then vortexed for 4 hours at room temp. The particles were then centrifugally precipitated using acetone and rinsed 3× with a 1:1 mixture of acetone to 75 mM TMAOH. See SI S6 for complete details.

### Covalent attachment to thermolysin (Conjugation)

A novel conjugation procedure was developed to get around the incompatibility of the solubility of the nanoparticles and thermolysin, which is soluble in high ionic strength solutions. First 100 mg of thermolysin (lyophilized type X, Sigma Aldrich) was desalted and buffer exchanged to 3 mL pH 7.5, 50 mM phosphate buffer. Then 10 mg of lyophilized and thoroughly rinsed Fe_3_O_4_@Au NP were dissolved in 3 mL fresh 50 mM pH 6.8 phosphate buffer. Then 10 mg of HATU was added to the nanoparticle solution and sonicated for 20 minutes. After sonication the HATU activated Au-NP solution was diluted with 6 mL of pH 6.8, 50 mM phosphate buffer and added directly to the thermolysin pellet, and lightly vortexed for 3 hours at 4°C. The conjugates were then precipitated via centrifuge, rinsed 2× and then re-suspended in 10 mL of pH 7.5, 50 mM phosphate buffer. No enzymatic activity was detected in the second rinsing, indicating very little if any free thermolysin left in solution. Besides testing the activity of the magnetic conjugates, a SDS PAGE was used to confirm conjugation. See SI S6 for complete details.

### Thermolysin activity assay and bulk heating experiments

The assay was based on the commercially available Pierce™ Colorimetric Protease Assay Kit (catalog # 23263) optimized for the specific enzyme conjugates and substrate. Aliquots (for 0,1,2,3 minute time points) of 50 μL of 4 mg/mL succinylated casein in 50 mM pH 7.5 50 mM phosphate buffer were pre-‘heated’ at 17.7°C (or different temperature for the temperature studies) in a thermocycler for ~10 minutes along with a buffer blank. Then 5 μL of thermolysin conjugates were added and the samples were heated for 0, 1, 2 and 3 minute time points. To stop the reaction 10 μL of 0.5 M EDTA was added to chelate the catalytic zinc and structural calcium ion, denaturing thermolysin. Then after allowing the reaction to cool back down to room temperature for 5 minutes 25 μL of colorimetric reagent, TNBSA was added. The samples were then incubated at room temperature for 20 minutes, and using the pedestal on a NanoDrop uv-Vis spectrometer the absorbance was measured at 430 nm to monitor formation of the orange product that forms when primary amines react with TNBSA. The slope of the line was then used to calculate the velocity of the enzyme reaction. See SI S9 for complete details. The same activity assay was used to determine kinetic parameters of the reaction by varying the substrate concentration. See SI S15 for complete details

### Radiofrequency heating assays

The exact same assay procedure was used for the RF heating experiments so that they were comparable. The fields were generated using a homemade water-cooled solenoid, with 14 turns and a length of 16 cm. We measured the field strength with a hall effect probe and found it was 0.021T in the center where the samples were placed. It was powered by a Philips PM5192 function generator to input a 17.76 MHz sinusoidal signal into a IFI sc×100 power amplifier. An impedance matching box was used to tune the reflected power to zero, new tunings were required for changes in frequency and applied power. See SI S12 for complete details.The bulk solution temperature was monitored with a Neoptix Nomad fiber optic temperature probe and then the conjugates were pre-heated in a thermocycler as described above to the same temperature, and the velocities of the two different heating methods were compared. Temperature measurements were made consistent by using a stand that holds the fiber optic probe in the same location for every sample. See SI S14 for complete details.

## Results and Discussion

We investigated the activity of Thermolysin covalently conjugated to 6.1 nm diameter gold coated magnetite particles under RF irradiation. Thermolysin is a 34.6 kDa metalloprotease from thermophilic bacteria Bacillus thermoproteolyticus [28–30]. It is active in the hydrolysis of peptide bonds with a preference for bonds between hydrophobic residues. Its activity is maximized at pH 8 and increases with temperature until 70°C at which point it denatures.

Magnetite particles were used because they are the superparamagnetic particles most frequently used in the literature for RF heating experiments. We gold coated the particles to stabilize the particles and allow for a robust covalent attachment of the conjugates through gold thiolate bonds, without negatively impacting the magnetic properties[31–33]. The optimal size magnetite cores were found for our system (17.7 MHz) to be 4 nm diameter [20]. We then coated the particles with the minimal amount of gold that would still allow for phase transferring and ligand exchange while maintaining solution stability. For complete details of the synthesis and phase transfer see SI page S3.

Thermolysin was attached to the gold coated iron-oxide particles through peptide bond formation using the carboxylic acid terminated ligands on the particles and the solvent accessible free lysine residues on thermolysin [34,35]. To significantly improve yield and decrease reaction time, a novel procedure was developed using HATU as a coupling reagent in buffer without any organic solvents (for details see SI page S5). This coupling procedure can be universally applied to enzymes, including those with very poor water solubility, a weak affinity for the ligand coating of the particle and high sensitivity to organic solvents like thermolysin.

The 17.76 MHz RF field was generated using a homemade water-cooled copper solenoid. A function generator was used to make a sinusoidal 17.76 MHz signal that was amplified and sent to the coil. An oscilloscope was used to monitor the output of the amplifier and an impedance matching box was used to ensure that the applied power was not reflected back into the power amplifier, see figure 1 for an illustration of the heating system. The temperature of the solution was measured using a fiber optic temperature probe. The water cooling system was used because it maintained a much more constant sample temperature than a thermal insulator without attenuating the RF field, across a wide range of applied powers. For complete system details see SI page S12.

**Figure 1.**
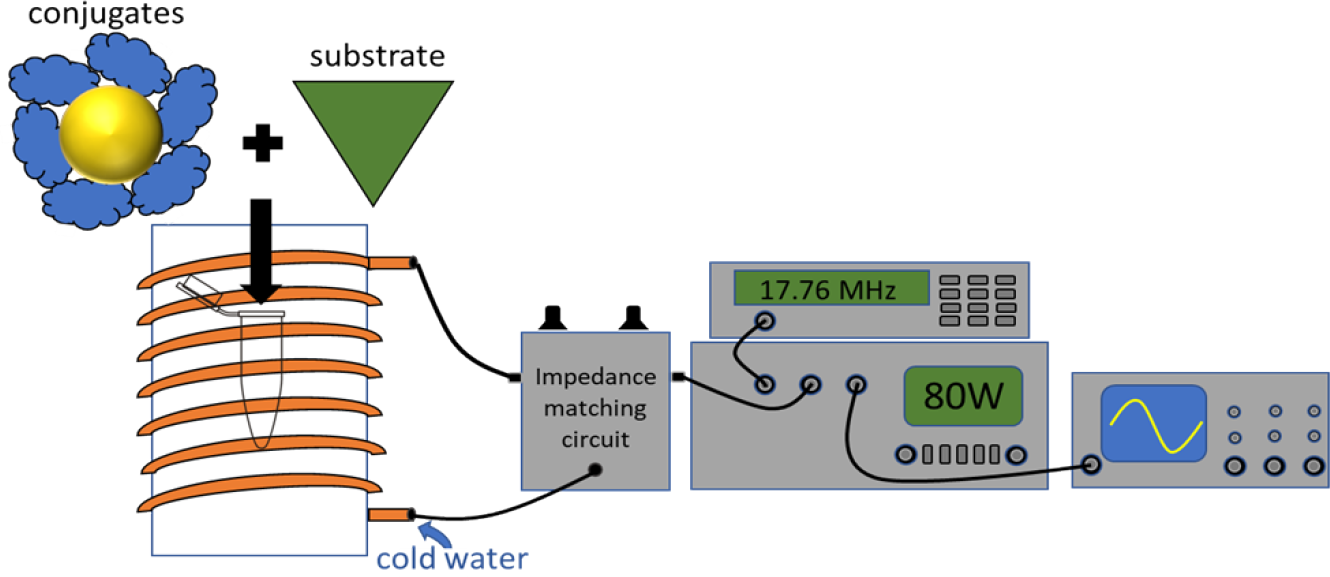
Illustration of radiofrequency heating system. A function generator is used to create a sinusoidal 17.76 MHz signal which is sent to a power amplifier. The signal is then sent through an impedance matching circuit and into the water cooled solenoid. For full details see SI page S12.

Preliminary experiments suggested higher power and higher frequency both increased the activity of enzyme/particle conjugates. The reported experiments were executed at 17.76 MHz and 80 W incident power, as these were the highest values of incident power and frequency that we could accomplish with our equipment.

Figure 2, left panel, illustrates the increase in observed thermolysin/NP activity by colorimetric assay that occurs under these RF conditions. Overall, we observe that the thermolsyin/NP conjugate activity is increased by 129 ± 8% under the 80 W, 17.76MHz RF irradiation when compared to a control reaction at the same solution temperature (as measured with a fiber optic (i.e.., metal-free) temperature probe). Figure 2, right panel shows the activity of each component of the thermolysin/NP conjugates (i.e., thermolsyin alone, NP alone) as well as the activity of thermolysin in the presence of unconjugated NP. Overall, we see a clear increase in activity of the enzyme relative to the controls. The measured maximum solution temperature in the RF fields was 17°C, indicating that the increase in enzyme activity observed is a local effect, and does not extend to heating the bulk solution. The modest activity increase of free thermolysin under RF irradiation is likely due to local regions of heat generated by ions oscillating with the electric component of the RF field. We do not attribute any activation of thermolysin to adsorption on the particles surface because thermolysin is very hydrophobic and our attempts to conjugate by simple adsorption were very unsuccessful. Overall, we interpret this set of control experiments as validating that the observed increase in activity under RF is attributable primarily to local temperature increase around the nanoparticle that increases the effective operating temperature of covalently attached enzymes.

The data in Figure 2 were all collected at 17°C solution temperature, which is far from the optimal thermolysin temperature of 70°C. By comparing the activity of thermolysin at various RF incident power values to the activity of thermolysin at various solution temperatures, we can propose effective ‘local’ temperature values for RF heated enzymes.

Figure 3 plots the activity of thermolysin as a function of solution temperature. We observe a linear relationship between temperature and enzyme rate (k), from 10°C to 70°C. At 80°C, the enzyme rate decreases, which we attribute to partial thermal denaturation (Figure 3, black triangles). We observe a strikingly similar relationship for activity as a function of incident power (Figure 3, red circles). At 80 W the in 17°C solution temperature conjugates perform as if they are in ~16°C warmer solution temperature.

**Figure 3.**
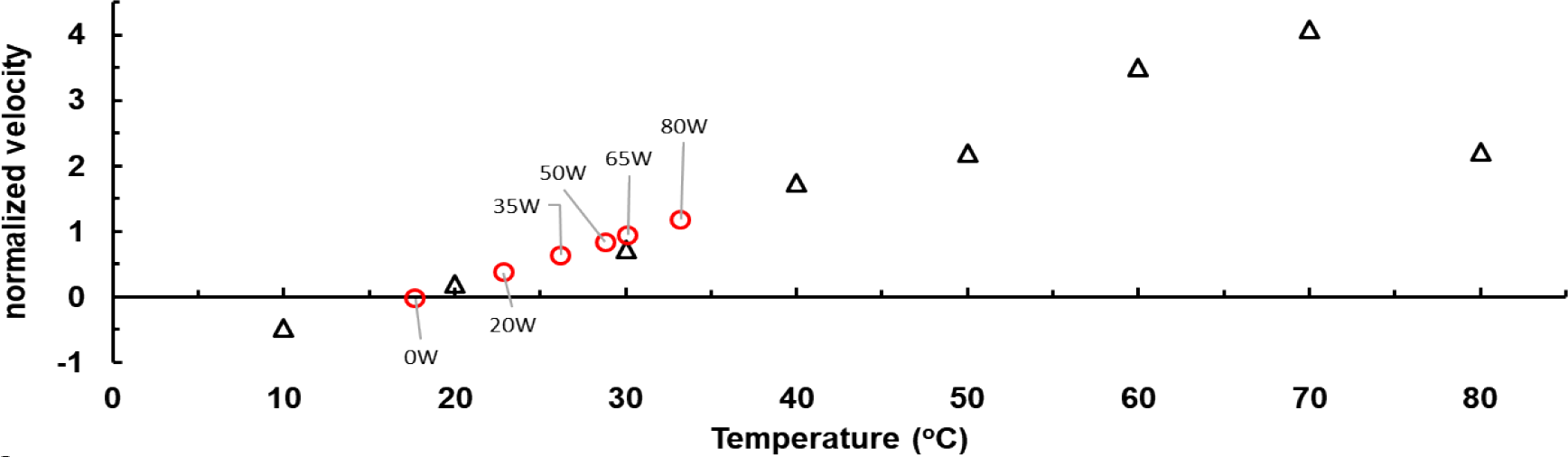
Plot of normalized activity of thermolysin as a function of temperature (black triangles) vs applied power (red circles). The zero point on the activity axis was set to the temperature of the solenoid without an RF field on.

We further investigated the enzyme kinetics of RF irradiated thermolysin/NP conjugates with concurrent controls identical in experimental setup except for the RF irradiation. Velocities were measured for various substrate concentrations. The results were analyzed with the Michaelis-Menten model of enzyme kinetics [36–39]. Figure 4 shows a double-reciprocal (i.e. Lineweaver-Burke plot, see SI page S15 for raw data). Analysis through equation 1 of this data produces the values for maximum enzyme velocity (V_max_), Michaelis constant (K_M_), and K_cat_. The two linear regressions were compared using GraphPad Prism 7.4 and with at a 95% confidence interval the regression coefficients are statistically different.

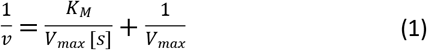

**Figure 4.**
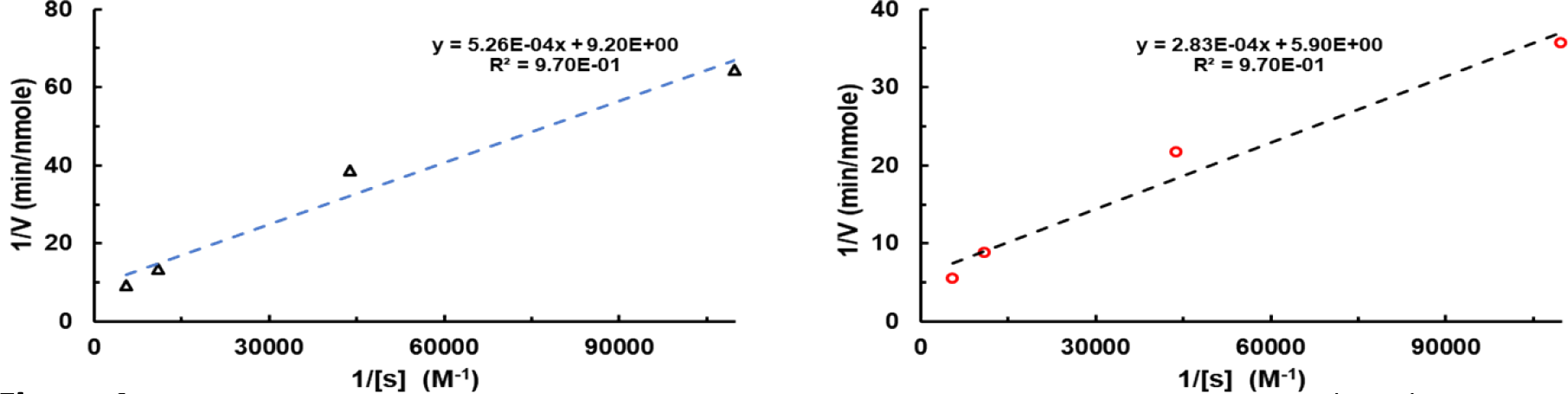
Double reciprocal plot of kinetics data, RF heated sample at 50W is in red circles (right), without RF at the same temperature is the black triangles (left). From the intercept and slope we determined K_m_ and V_max_ and k_cat_ using the Michaelis-Menten model of enzyme kinetics.

These values are tabulated in Table 1. Under RF irradiation, the V_max_ of thermolsyin/NP conjugates increase relative to the same sample in the absence of RF. We rationalize this by considering that transition states will be more stable (higher thermal energy results in lower activation energy barriers which speeds any reaction up [40]. More curiously, we observe a decrease in K_M_ in RF fields. This may be understood through an examination of the thermolysin mechanism. Thermolysin most likely utilizes an induced fit mechanism of activity in which substrate molecules induces conformational changes in thermolysin that help it stabilize transition states [41]. Under typical heating conditions, global temperature increases the floppiness of the enzyme and substrate resulting in a smearing of states of the enzyme and substrate from the ideal state, but allows for more sampling of states.

**Table 1.** Summary of kinetics data for Michaelis Menten kinetics model.

|  | Radiofrequency | Bulk Control |
| --- | --- | --- |
| $V_{max}$ (nmole/min) | $0.169 \pm 0.059$ | $0.109 \pm 0.044$ |
| $K_m$ ($\mu M$) | $48.0 \pm 6.0$ | $57.2 \pm 6.8$ |
| $k_{cat}$ ($min^{-1}$) | $2.1 \cdot 10^{-4}$ | $1.4 \cdot 10^{-4}$ |

The decrease in K_M_ (i.e., increase in stability of intermediate substrate enzyme complex) that occurs as a result of RF heating can be rationalized in terms of enzymatic mechanism. Thermolysin active site structure is modified by the substrate – an induced fit [41,42]. The substrate induced conformational changes in thermolysin structure stabilize the proteolytic transition state. Higher temperatures increase the rate of conformational sampling that the enzyme does in making an induced fit [40,43]. However, higher temperatures also increase the conformational sampling of the substrate, making induced fit less likely. In terms of the Michaelis-Menten model (scheme 1), higher temperatures increase both k_1_ and k_−1_. In bulk heating experiments K_M_ increases with temperature because increased temperatures increase k_cat_ and influence enzyme substrate binding forward and backward rates equally.

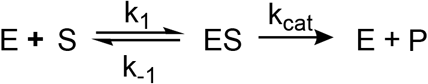

**Scheme 1.** Simplified enzymatic reaction for Michaelis Menten kinetics model. Its

In the radio-frequency fields, the enzyme is closer to the heat source than the substrate. Thus, the enzyme is sampling conformational space at a faster rate than the more rigid substrate. We suggest that this situation of ‘hot enzyme, cold substrate’ allows for a more rapid induced fit the enzyme is being heated results in additional floppiness of the enzyme, allowing for an easier induced fit of the substrate molecule. This results in higher k_1_ because the substrate is docking with a floppier enzyme (allowing for best confirmation more easily) and then lower k_−1_ because the substrate is held more firmly once its attached because of this same improvement in induced fit through enzyme floppiness [40].

## Conclusions

We have successfully shown that thermolysin can be remotely activated in 17.76 MHz RF fields when covalently attached to magnetic nanoparticles. Without raising the bulk solution temperature, under RF irradiation we observed enzyme activity as if the solution was 16 ± 2 °C warmer, or an increase in enzymatic rate of 129 ± 8%. Further studies will be performed with higher field strengths to maximize the level of activation in RF fields. A preliminary investigation of the kinetics in and outside of the RF fields showed an increase in activity consistent with a hot enzyme interacting with a cold substrate. Our approach for remote control of enzyme activity seems to be universally applicable to any enzyme that has an induced fit mechanism of activity, but more investigation into the scope of this process is necessary.

## Acknowledgements

This work was supported by NIH R01 GM112225.

